# Monod-Wyman-Changeux Analysis of Ligand-Gated Ion Channel Mutants

**DOI:** 10.1101/102194

**Authors:** Tal Einav, Rob Phillips

**Affiliations:** Department of Physics, California Institute of Technology, Pasadena, California 91125, United States; Department of Applied Physics and Division of Biology and Biological Engineering, California Institute of Technology, Pasadena, California 91125, United States

## Abstract

We present a framework for computing the gating properties of ligand-gated ion channel mutants using the Monod-Wyman-Changeux (MWC) model of allostery. We derive simple analytic formulas for key functional properties such as the leakiness, dynamic range, half-maximal effective concentration ([*EC*_50_]), and effective Hill coefficient, and explore the full spectrum of phenotypes that are accessible through mutations. Specifically, we consider mutations in the channel pore of nicotinic acetylcholine receptor (nAChR) and the ligand binding domain of a cyclic nucleotide-gated (CNG) ion channel, demonstrating how each mutation can be characterized as only affecting a subset of the biophysical parameters. In addition, we show how the unifying perspective offered by the MWC model allows us, perhaps surprisingly, to collapse the plethora of dose-response data from different classes of ion channels into a universal family of curves.

## 1 Introduction

Ion channels are signaling proteins responsible for a huge variety of physiological functions ranging from responding to membrane voltage, tension, and temperature to serving as the primary players in the signal transduction we experience as vision.^1^ Broadly speaking, these channels are classified based upon the driving forces that gate them. In this work, we explore one such classification for ligand-gated ion channel mutants based on the Monod-Wyman-Changeux (MWC) model of allostery. In particular, we focus on mutants in two of the arguably best studied ligand-gated ion channels: the nicotinic acetylcholine receptor (nAChR) and the cyclic nucleotide-gated (CNG) ion channel shown schematically in Fig.1.^2,3^

**Figure 1:**
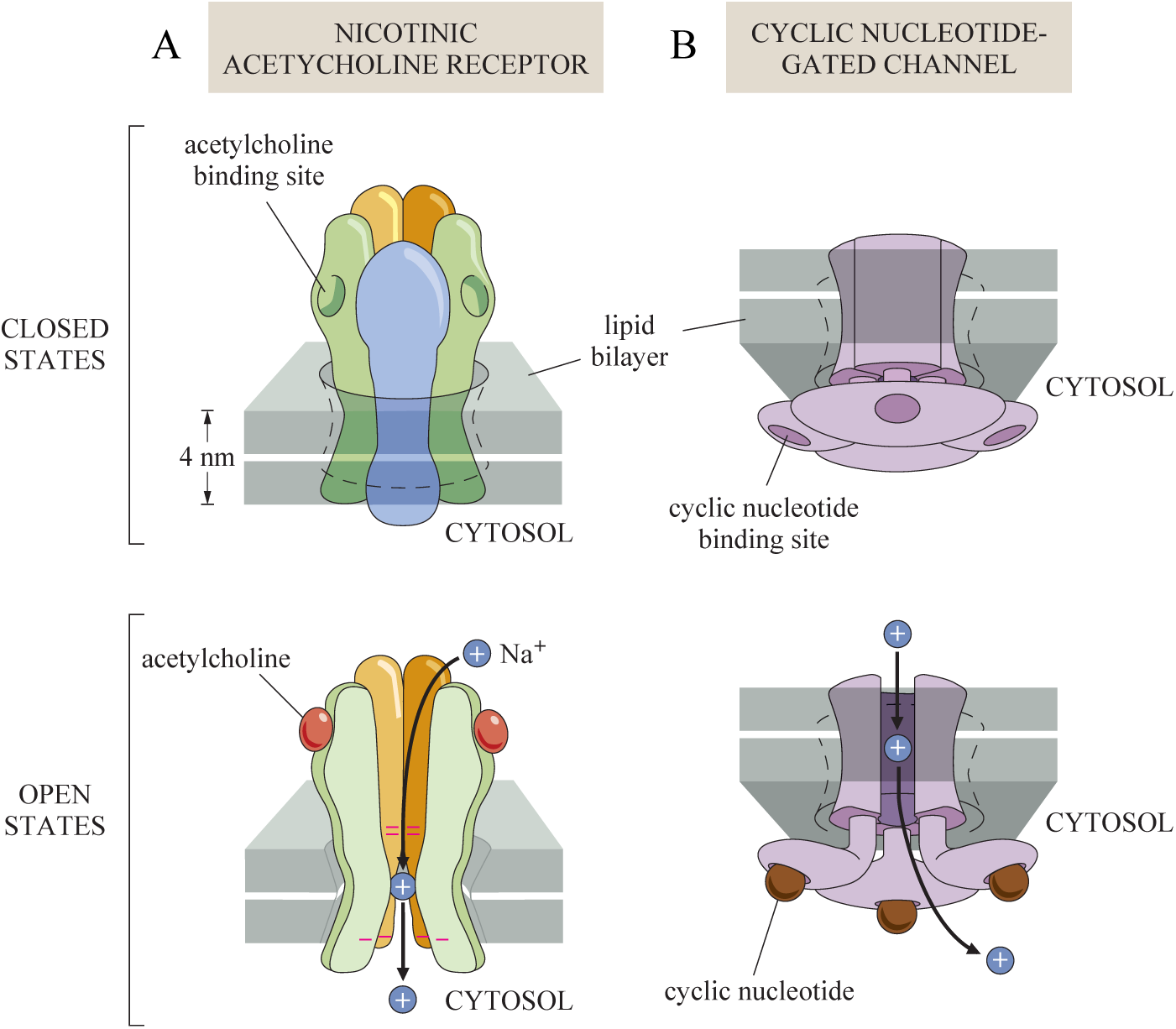
Schematic of nAChR and CNGA2 ion channels. (A) The heteropentameric nicotinic acetylcholine receptor (nAChR) has two ligand binding sites for acetylcholine outside the cytosol. (B) The homotetrameric cyclic nucleotide-gated (CNGA2) has four ligand binding sites, one on each subunit, for cAMP or cGMP located inside the cytosol. Both ion channels have a higher probability of being closed in the absence of ligand and open when bound to ligand.

The MWC model has long been used in the contexts of both nAChR and CNG ion channels.^4–6^ Although careful analysis of these systems has revealed that some details of ligand-gated ion channel dynamics are not captured by this model (e.g. the existence and interaction of multiple conducting states^7,8^), the MWC framework nevertheless captures many critical features of a channel’s response and offers one of the simplest settings to explore its underlying mechanisms. For example, knowledge of both the nAChR and CNG systems’ molecular architecture and our ability to measure their complex kinetics has only recently become sufficiently advanced to tie the effects of mutations to key biophysical parameters. Purohit and Auerbach used combinations of mutations to infer the nAChR gating energy, finding that unliganded nAChR channels open up for a remarkably brief 80 *μ*s every 15 minutes.^9^ Statistical mechanics has been used to show how changes to the energy difference between conformations in allosteric proteins translates to different functional behavior (i.e. how it modifies the leakiness, dynamic range, [*EC*_50_] and the effective Hill coefficient),^10,11^ and we extend this work to find simple analytic approximations that are valid within the context of ion channels. Using this methodology, we systematically explore the full range of behaviors that may be induced by different types of mutations. This analysis enables us to quantify the inherent trade-offs between key properties of ion channel dose-response curves and potentially paves the way for future biophysical models of evolutionary fitness in which the genotype (i.e. amino acid sequence) of allosteric molecules is directly connected to phenotype (i.e. properties of a channel’s response).

To this end, we consider two distinct classes of mutants which tune different sets of MWC parameters – either the ion channel gating energy or the ligand-channel dissociation constants. Previous work by Auerbach *et al*. demonstrated that these two sets of physical parameters can be independently tuned within the nAChR ion channel; pore mutations only alter the channel gating energy whereas mutations within the ligand binding domain only affect the ligand-channel dissociation constants.^12^ Utilizing this parameter independence, we determine the full spectrum of nAChR phenotypes given an arbitrary set of channel pore mutations and show why a linear increase in the channel gating energy leads to a logarithmic shift in the nAChR dose-response curve. Next, we consider recent data from CNGA2 ion channels with mutations in their ligand binding pocket.^13^ We hypothesize that modifying the ligand binding domain should not alter the channel gating energy and demonstrate how the entire class of CNGA2 mutants can be simultaneously characterized with this constraint. This class of mutants sheds light on the fundamental differences between homooligomeric channels comprised of a single type of subunit and heterooligomeric channels whose distinct subunits can have different ligand binding affinities.

By viewing mutant data through its effects on the underlying biophysical parameters of the system, we go well beyond simply fitting individual dose-response data, instead creating a framework with which we can explore the full expanse of ion channel phenotypes available through mutations. Using this methodology, we: (1) analytically compute important ion channel characteristics, namely the leakiness, dynamic range, [*EC*_50_], and effective Hill coefficient, (2) link the role of mutations with thermodynamic parameters, (3) show how the behavior of an entire family of mutants can be predicted using only a subset of the members of that family, (4) quantify the pleiotropic effect of point mutations on multiple phenotypic traits and characterize the correlations between these diverse effects, and (5) collapse the data from multiple ion channels onto a single master curve, revealing that such mutants form a one-parameter family. In doing so, we present a unified framework to collate the plethora of data known about such channels.

## Model

Electrophysiological techniques can measure currents across a single cell’s membrane. The current flowing through a ligand-gated ion channel is proportional to the average probability *p*_open_(*c*) that the channel will be open at a ligand concentration *c*. For an ion channel with *m* identical ligand binding sites (see Fig 2), this probability is given by the MWC model as

**Figure 2:**
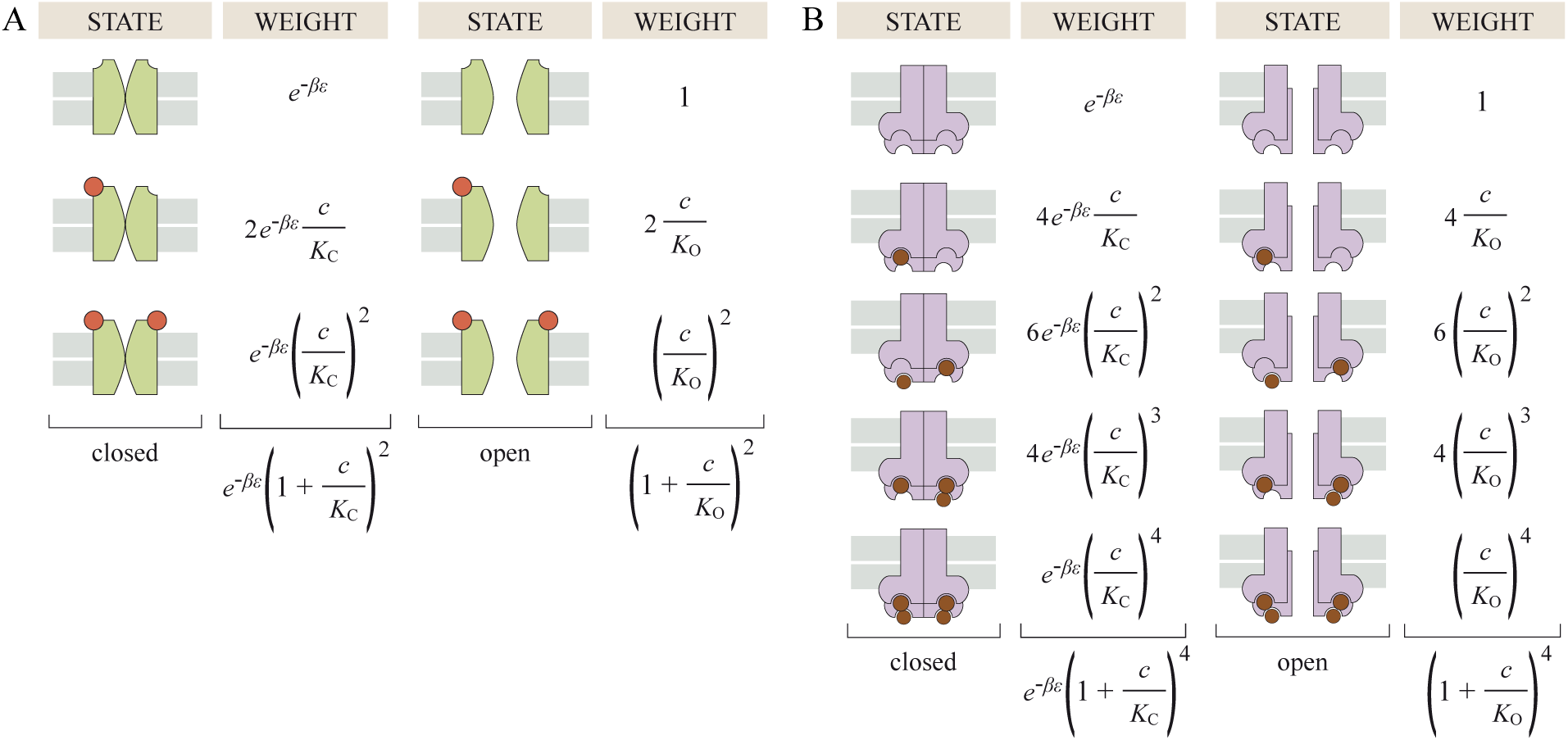
The probability that a ligand-gated ion channel is open as given by the MWC model. (A) Microscopic states and Boltzmann weights of the nAChR ion channel (green) binding to acetylcholine (orange). (B) Corresponding states for the CNGA2 ion channel (purple) binding to cGMP (brown). The behavior of these channels is determined by three physical parameters: the affinity between the receptor and ligand in the open (*K*_O_) and closed (*K*_C_) states and the free energy difference between the closed and open conformations of the ion channel.

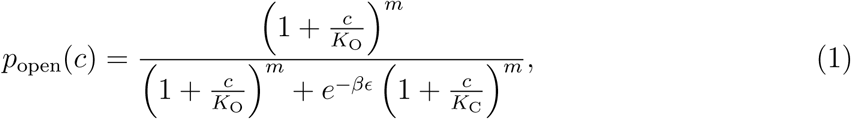

where *K*_O_ and *K*_C_ represent the dissociation constants between the ligand and the open and closed ion channel, respectively, *c* denotes the concentration of the ligand, *ϵ* (called the gating energy) denotes the free energy difference between the closed and open conformations of the ion channel in the absence of ligand, and 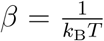 where *k*_B_ is Boltzmann’s constant and *T* is the temperature of the system. Wild type ion channels are typically closed in the absence of ligand (*ϵ < 0*) and open when bound to ligand (*K*_O_ < *K*_C_). Fig 2 shows the possible conformations of the nAChR (*m* = 2) and CNGA2 (*m* = 4) ion channels together with their Boltzmann weights. *p*_open_(*c*) is given by the sum of the open state weights divided by the sum of all weights. Note that the MWC parameters *K*_O_, *K*_C_, and *ϵ* may be expressed as ratios of the experimentally measured rates of ligand binding and unbinding as well as the transition rates between the open and closed channel conformations (see Supporting Information section A.1).

Current measurements are often reported as *normalized* current, implying that the current has been stretched vertically to run from 0 to 1, as given by

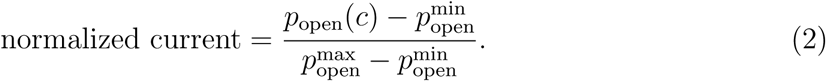

*p*_open_(*c*) increases monotonically as a function of ligand concentration *c*, with a minimum value in the absence of ligand given by

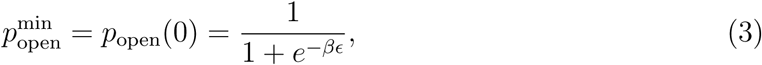

and a maximum value in the presence of saturating levels of ligand given as

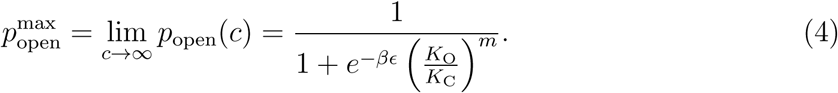

Using the above two limits, we can investigate four important characteristics of ion channels.^10,11^ First, we examine the *leakiness* of an ion channel, or the fraction of time a channel is open in the absence of ligand, namely,

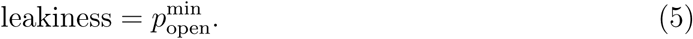

Next we determine the *dynamic range*, or the difference between the probability of the maximally open and maximally closed states of the ion channel, given by

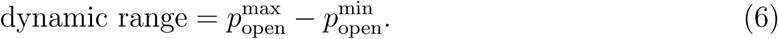

Ion channels that minimize leakiness only open upon ligand binding, and ion channels that maximize dynamic range have greater contrast between their open and closed states. Just like *p*_open_(*c*), leakiness and dynamic range lie within the interval [0; 1].

Two other important characteristics are measured from the normalized current. The *half maximal effective concentration* [*EC*_50_] denotes the concentration of ligand at which the normalized current of the ion channel equals ½, namely,

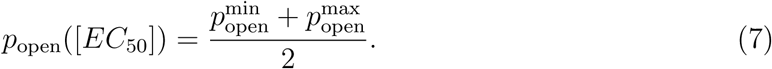

The *effective Hill coefficient h* equals twice the log-log slope of the normalized current evaluated at *c* = [*EC*_50_],

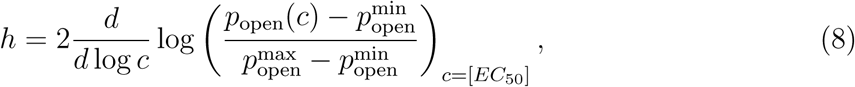

Which reduces to the standard Hill coefficient for the Hill function.^14^ The [*EC*_50_] determines how the normalized current shifts left and right, while the effective Hill coefficient corresponds to the slope at [*EC*_50_]. Together, these two properties determine the approximate window of ligand concentrations for which the normalized current transitions from 0 to 1.

In the limit 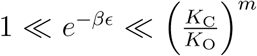, which we show below is relevant for both the nAChR and CNGA2 ion channels, the various functional properties of the channel described above can be approximated to leading order as (see Supporting Information section B):

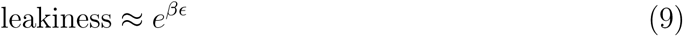

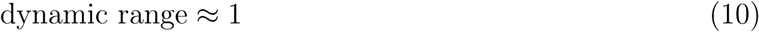

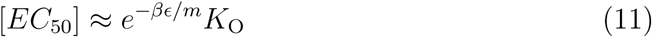

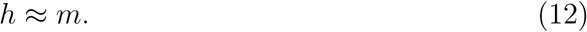

## Results

### nAChR Mutants can be Categorized using Free Energy

Muscle-type nAChR is a heteropentamer with subunit stoichiometry *α*_2_*βγδ*, containing two ligand binding sites for acetylcholine at the interface of the *α-δ* and *α-γ* subunits.^15^ The five homologous subunits have M2 transmembrane domains which move symmetrically during nAChR gating to either occlude or open the ion channel.^16^ By introducing a serine in place of the leucine at a key residue (L251S) within the M2 domain present within each subunit, the corresponding subunit is able to more easily transition from the closed to open configuration, shifting the dose-response curve to the left (see Fig 3A).^17^ For example, wild type nAChR is maximally stimulated with 100 *μ*M of acetylcholine, while a mutant ion channel with one L251S mutation is more sensitive and only requires 10 *μ*M to saturate its dose-response curve.

**Figure 3:**
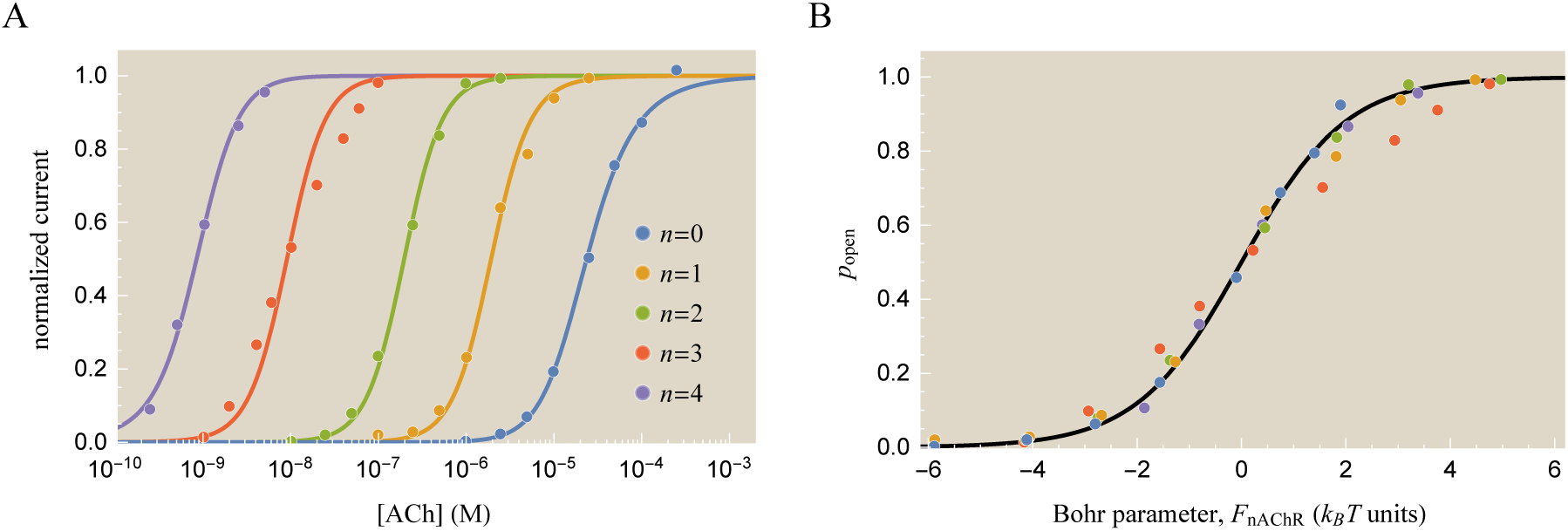
Characterizing nicotinic acetylcholine receptors with *n* subunits carrying the L251S mutation. (A) Normalized currents of mutant nAChR ion channels at different concentrations of the agonist acetylcholine (ACh).^17^ The curves from right to left show a receptor with *n* = 0 (wild type), *n* = 1 (*α*_2_*βγ*\**δ*), 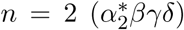, *n* = 3 (*α*_2_*β*\**γ*\**δ*\*), and 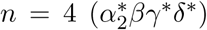 mutations, where asterisks (*) denote a mutated subunit. Fitting the data (solid lines) to Eqs (1) and (2) with *m* = 2 ligand binding sites determines the three MWC parameters *K*_O_ = 0.1 × 10^−9^ M, *K*_C_ = 60 × 10^−6^ M, and *βϵ*^(*n*)^ = [−4.0, −8.5, −14.6, −19.2, −23.7] from left (*n* = 4) to right (*n* = 0). With each additional mutation, the dose-response curve shifts to the left by roughly a decade in concentration while the *ϵ* parameter increases by roughly 5 *k*_B_*T*. (B) The probability *p*_open_(*c*) that the five ion channels are open can be collapsed onto the same curve using the Bohr parameter *F*_nAChR_(*c*) given by Eq (13). A positive Bohr parameter indicates that *c* is above the [*EC*_50_]. See Supporting Information section C for details on the fitting procedure.

Labarca *et al.* used L251S mutations to create ion channels with *n* mutated subunits.^17^ Fig 3A shows the resulting normalized current for several of these mutants; from right to left the curves represent *n* = 0 (wild type) to *n* = 4 (an ion channel with four of its five subunits mutated). One interesting trend in the data is that each additional mutation shifts the normalized current to the left by approximately one decade in concentration (see Supporting Information section A.2). This constant shift in the dose-response curves motivated Labarca *et al.* to postulate that mutating each subunit increases the gating free energy *ϵ* by a fixed amount.

To test this idea, we analyze the nAChR data at various concentrations *c* of the ligand acetylcholine using the MWC model Eq (1) with *m* = 2 ligand binding sites. Because the L251S mutation is approximately 4:5 nm from the ligand binding domain,^18^ we assume that the ligand binding affinities *K*_O_ and *K*_C_ are unchanged for the wild type and mutant ion channels, an assumption that has been repeatedly verified by Auerbach *et al.* for nAChR pore mutations.^12^ Fig 3A shows the best-fit theoretical curves assuming all five nAChR mutants have the same *K*_O_ and *K*_C_ values but that each channel has a distinct gating energy *ϵ*^(*n*)^ (where the superscript *n* denotes the number of mutated subunits). These gating energies were found to increase by roughly 5 *k_B_T* per *n*, as would be expected for a mutation that acts equivalently and independently on each subunit.

One beautiful illustration of the power of the MWC model lies in its ability to provide a unified perspective to view data from many different ion channels. Following earlier work in the context of both chemotaxis and quorum sensing,^19,20^ we rewrite the probability that the nAChR receptor is open as

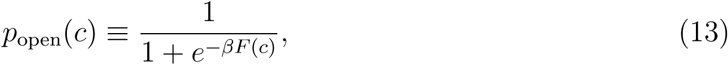

where this last equation defines the *Bohr parameter*^21^

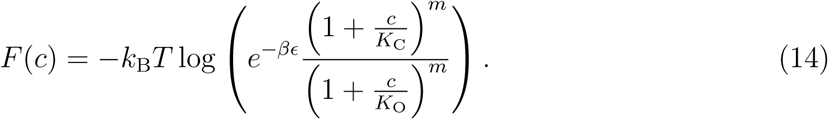

The Bohr parameter quantifies the trade-offs between the physical parameters of the system (in the case of nAChR, between the entropy associated with the ligand concentration *c* and the gating free energy *βϵ*). When the Bohr parameters of two ion channels are equal, both channels will elicit the same physiological response. Using Eqs (1) and (13) to convert the normalized current data into the probability *p*_open_ (see Supporting Information section A.3), we can collapse the dose-response data of the five nAChR mutants onto a single master curve as a function of the Bohr parameter for nAChR, *F*_nAChR_(*c*), as shown in Fig 3B. In this way, the Bohr parameter maps the full complexity of a generic ion channel response into a single combination of the relevant physical parameters of the system.

### Full Spectrum of nAChR Gating Energy Mutants

We next consider the entire range of nAChR phenotypes achievable by only modifying the gating free energy *ϵ* of the wild type ion channel. For instance, any combination of nAChR pore mutations would be expected to not affect the ligand dissociation constants and thus yield an ion channel within this class (see Supporting Information section A.4 for one such example). For concreteness, we focus on how the *ϵ* parameter tunes key features of the dose-response curves, namely the leakiness, dynamic range, [*EC*_50_], and effective Hill coefficient *h* (see Eqs (5)-(12)), although we note that other important phenotypic properties such as the intrinsic noise and capacity have also been characterized for the MWC model.^10^ Fig 4 shows these four characteristics, with the open squares representing the properties of the five best-fit dose-response curves from Fig 3A.

**Figure 4:**
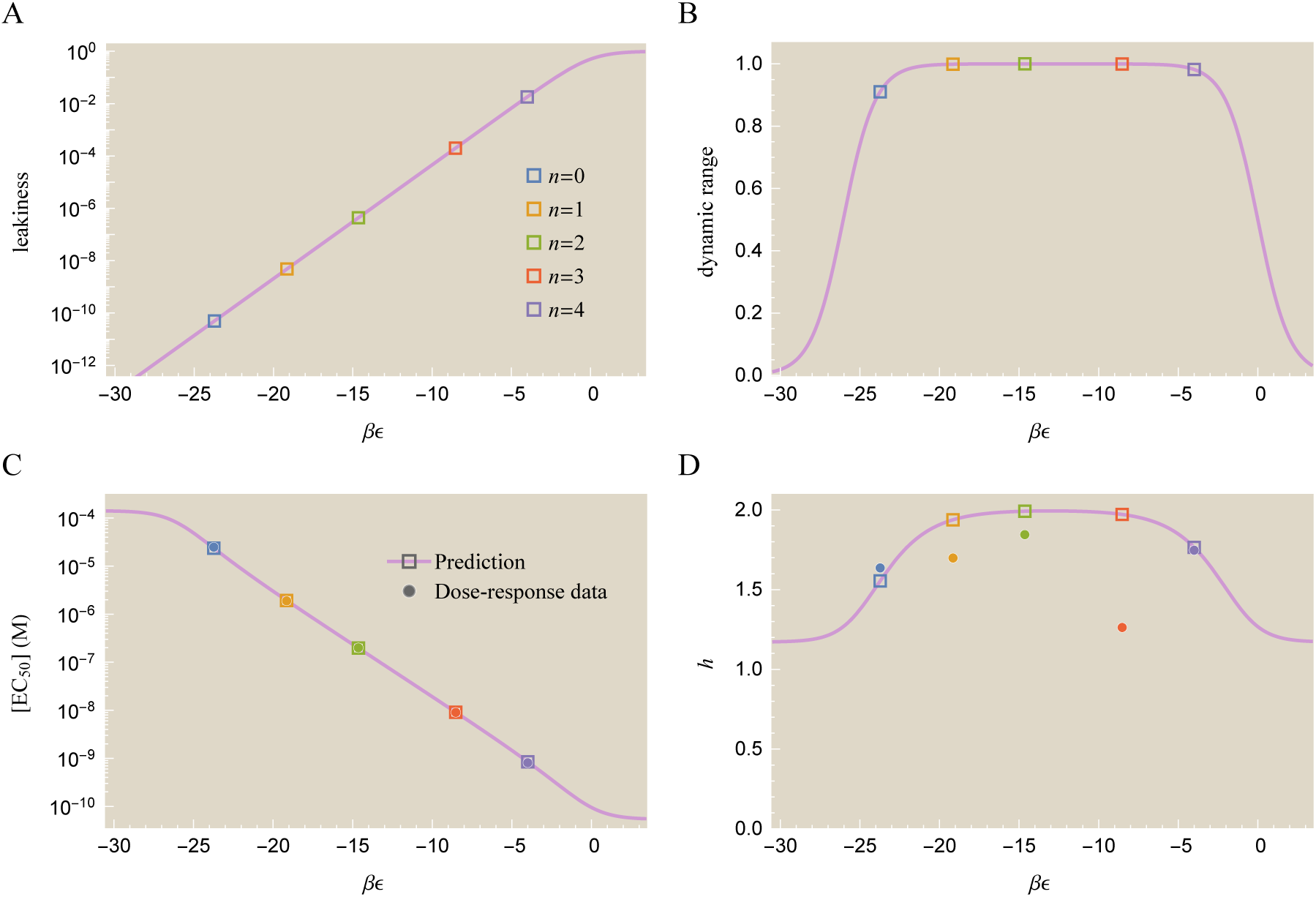
Theoretical prediction and experimental measurements for mutant nAChR ion channel characteristics. The open squares mark the *βϵ* values of the five dose response curves from Fig 3A. (A) The leakiness given by Eq (5) increases exponentially with each mutation. (B) The dynamic range from Eq (6) is nearly uniform for all mutants. (C) The [*EC*_50_] decreases exponentially with each mutation. (D) The effective Hill coefficient *h* is predicted to remain approximately constant. [*EC*_50_] and *h* offer a direct comparison between the best-fit model predictions (open squares) and the experimental measurements (solid circles) from Fig 3A. While the [*EC*_50_] matches well between theory and experiment, the effective Hill coefficient *h* is significantly noisier.

Fig 4A implies that all of the mutants considered here have negligible leakiness; according to the MWC parameters found here, the probability that the wild type channel (*βϵ*^(0)^ = −23.7) will be open is less than 10^−10^. Experimental measurements have shown that such spontaneous openings occur extremely infrequently in nAChR,^22^ although direct measurement is difficult for such rare events. Other mutational analysis has predicted gating energies around *βϵ*^(0)^ ≈ −14 (corresponding to a leakiness of 10^−6^),^12^ but we note that such a large wild type gating energy prohibits the five mutants in Fig 3 from being fit as a single class of mutants with the same *K*_O_ and *K*_C_ values (see Supporting Information section C.2). If this large wild type gating energy is correct, it may imply that the L251S mutation also affects the *K*_O_ and *K*_C_ parameters, though the absence of error bars on the original data make it hard to quantitatively assess the underlying origins of these discrepancies.

Fig 4B asserts that all of the mutant ion channels should have full dynamic range except for the wild type channel, which has a dynamic range of 0.91. In comparison, the measured dynamic range of wild type nAChR is 0.95, close to our predicted value.^12^ Note that only when the dynamic range approaches unity does the normalized current become identical to *p*_open_; for lower values, information about the leakiness and dynamic range is lost by only measuring normalized currents.

We compare the [*EC*_50_] (Fig 4C) and effective Hill coefficient *h* (Fig 4D) with the nAChR data by interpolating the measurements (see Supporting Information section C.3) in order to precisely determine the midpoint and slope of the response. The [*EC*_50_] predictions faithfully match the data over four orders of magnitude. Because each additional mutation lowers the [*EC*_50_] by approximately one decade, the analytic form Eq (11) implies that *ϵ* increases by roughly 5 *k*_B_*T* per mutation, enabling the ion channel to open more easily. In addition to the L251S mutation considered here, another mutation (L251T) has also been found to shift [*EC*_50_] by a constant logarithmic amount (see Supporting Information section A.4).^23^ We also note that many biological systems logarithmically tune their responses by altering the energy difference between two allosteric states, as seen through processes such as phosphorylation and calmodulin binding.24 This may give rise to an interesting interplay between physiological time scales where such processes occur and evolutionary time scales where traits such as the [*EC*_50_] may be accessed via mutations like those considered here.^25^

Lastly, the Hill coefficients of the entire class of mutants all lie between 1.5 and 2.0 except for the *n* = 3 mutant whose dose-response curve in Fig 3A is seen to be flatter than the MWC prediction. We also note that if the L251S mutation moderately perturbs the *K*_O_ and *K*_C_ values, it would permit fits that more finely attune to the specific shape of each mutant’s data set. That said, the dose-response curve for the *n* = 3 mutant could easily be shifted by small changes in the measured values and hence without recourse to error bars, it is difficult to make definitive statements about the value adopted for *h* for this mutant.

Note that the simplified expressions Eqs (9)-(12) for the leakiness, dynamic range, [*EC*_50_], and effective Hill coefficient apply when 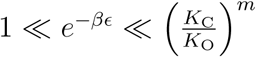, which given the values of *K*_C_ and *K*_C_ for the nAChR mutant class translates to 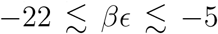. The *n* = 1, 2, and 3 mutants all fall within this range, and hence each subsequent mutation exponentially increases their leakiness and exponentially decreases their [*EC*_50_], while their dynamic range and effective Hill coefficient remain indifferent to the L251S mutation. The *βϵ* parameters of the *n* = 0 and *n* = 4 mutants lie at the edge of the region of validity, so higher order approximations can be used to more precisely fit their functional characteristics (see Supporting Information section B).

### Heterooligomeric CNGA2 Mutants can be Categorized using an Expanded MWC Model

The nAChR mutant class discussed above had two equivalent ligand binding sites, and only the gating free energy *βϵ* varied for the mutants we considered. In this section, we use beautiful data for the olfactory CNGA2 ion channel to explore the unique phenotypes that emerge from a heterooligomeric ion channel whose subunits have different ligand binding strengths.

The wild type CNGA2 ion channel is made up of four identical subunits, each with one binding site for the cyclic nucleotide ligands cAMP or cGMP.^26^ Within the MWC model, the probability that this channel is open is given by Eq (1) with *m* = 4 ligand binding sites (see Fig 2B). Wongsamitkul et al. constructed a mutated subunit with lower affinity for ligand and formed tetrameric CNGA2 channels from different combinations of mutated and wild type subunits (see Fig 5).^13^ Since the mutation specifically targeted the ligand binding sites, these mutant subunits were postulated to have new ligand dissociation constants but the same free energy difference *βϵ*.

**Figure 5:**
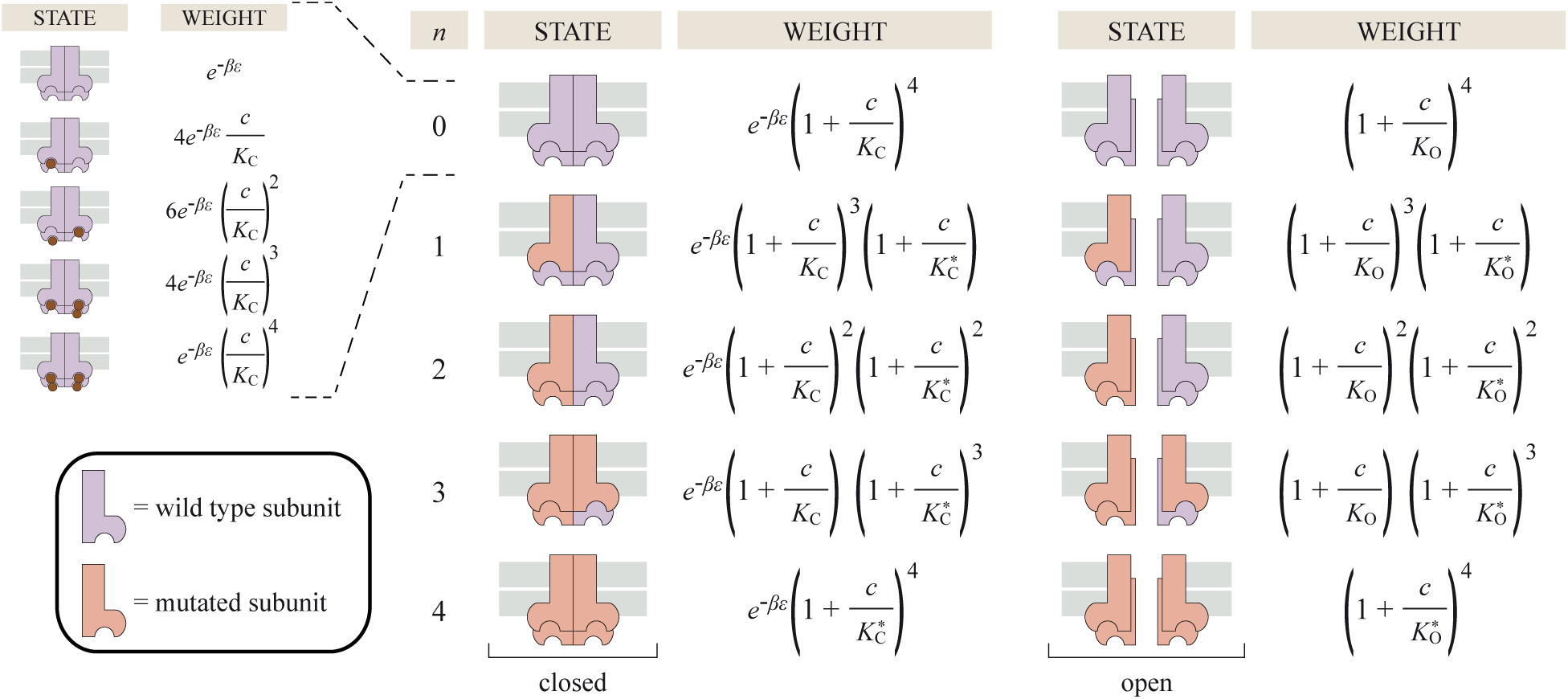
States and weights for mutant CNGA2 ion channels. CNGA2 mutants with *m* = 4 subunits were constructed using *n* mutated (light red) and *m* – *n* wild type subunits (purple). The affinity between the wild type subunits to ligand in the open and closed states (*K*_O_ and *K*_C_) is stronger than the affinity of the mutated subunits (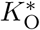 and 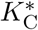). The weights shown account for all possible ligand configurations, with the inset explicitly showing all of the closed states for the wild type (*n* = 0) ion channel from Fig 2B. The probability that a receptor with *n* mutated subunits is open is given by its corresponding open state weight divided by the sum of open and closed weights in that same row.

We can extend the MWC model to compute the probability *p*_open_ that these CNGA2 constructs will be open. The states and weights of an ion channel with *n* mutated subunits (with ligand affinities 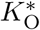 and 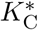) and *m* – *n* wild type subunits (with ligand affinities *K*_O_ and *K*_C_) is shown in Fig 5, and its probability to be open is given by

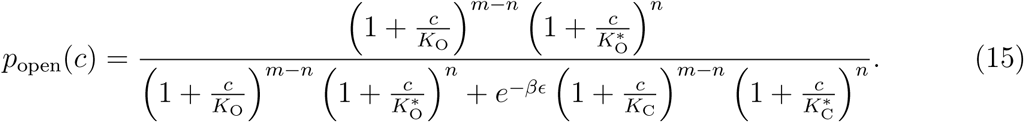

Measurements have confirmed that the dose-response curves of the mutant CNGA2 channels only depend on the total number of mutated subunits *n* and not on the positions of those subunits (for example both *n* = 2 with adjacent mutant subunits and *n* = 2 with mutant subunits on opposite corners have identical dose-response curves).^13^

Fig 6A shows the normalized current of all five CNGA2 constructs fit to a single set of *K*_O_, *K*_C_, 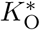, 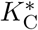, and *ϵ* parameters. Since the mutated subunits have weaker affinity to ligand (leading to the larger dissociation constants 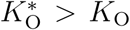 and 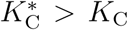), the [*EC*_50_] shifts to the right as *n* increases. As in the case of nAChR, we can collapse the data from this family of mutants onto a single master curve using the Bohr parameter *F*_CNGA2_(*c*) from Eqs (13) and (15), as shown in Fig 6B.

**Figure 6:**
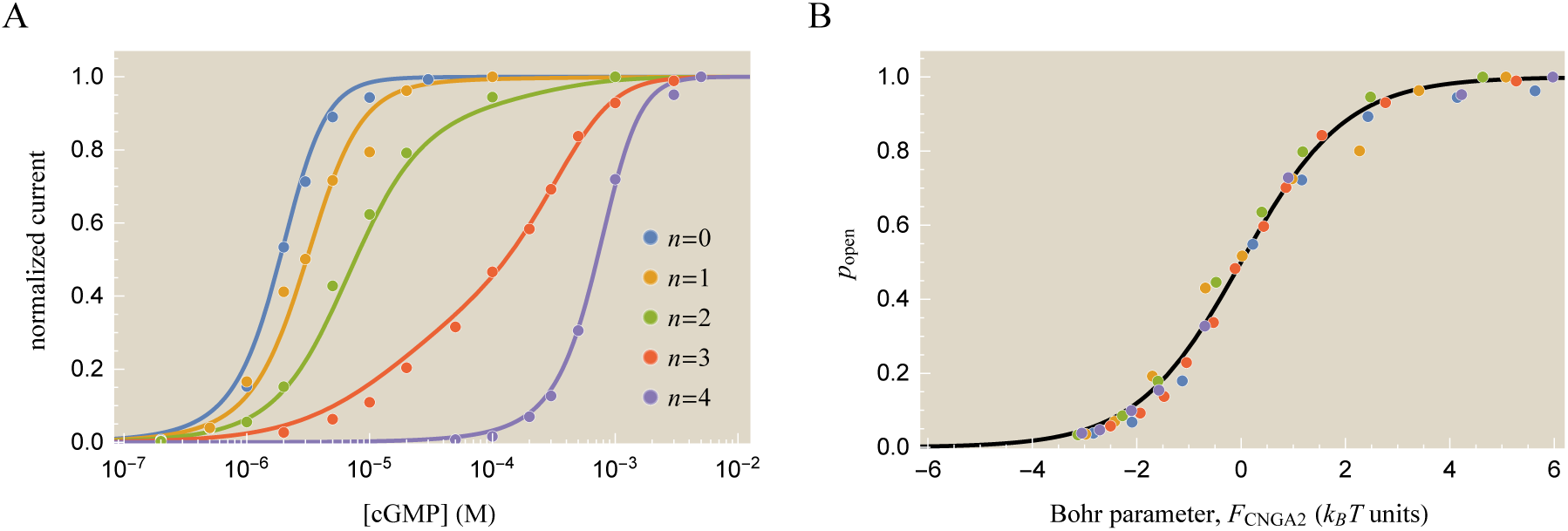
Normalized currents for CNGA2 ion channels with a varying number *n* of mutant subunits. (A) Dose-response curves for CNGA2 mutants comprised of 4 − *n* wild type subunits and *n* mutated subunits with weaker affinity for the ligand cGMP.^13^ Once the free energy *ϵ* and the ligand dissociation constants of the wild type subunits (*K*_O_ and *K*_C_) and mutated subunits (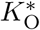 and 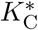) are fixed, each mutant is completely characterized by the number of mutated subunits *n* in Eq (15). Theoretical best-fit curves are shown using the parameters *K*_O_ = 1.2 × 10^−6^ M, *K*_C_ = 20 × 10^−6^ M, 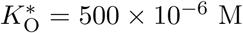, 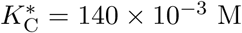, and *βϵ* = −3.4. (B) Data from all five mutants collapses onto a single master curve when plotted as a function of the Bohr parameter given by Eq (13). See Supporting Information section C for details on the fitting.

Although we analyze the CNGA2 ion channels in equilibrium, we can glimpse the dynamic
nature of the system by computing the probability of each channel conformation. Fig 7A shows the ten possible states of the wild type (*n* = 0) channel, the five open states *O_j_* and the five closed states *C_j_* with 0 ≤ *j* ≤ 4 ligands bound. Fig 7B shows how the probabilities of these states are all significantly shifted to the right in the fully mutated (*n* = 4) channel since the mutation diminishes the channel-ligand affinity. The individual state probabilities help determine which of the intermediary states can be ignored when modeling. One extreme simplification that is often made is to consider the Hill limit, where all of the states are ignored save for the closed, unbound ion channel (*C*_0_) and the open, fully bound ion channel (*O*_4_). The drawbacks of such an approximation are two-fold: (1) at intermediate ligand concentrations (*c* ∈ [10^−7^, 10^−5^]M for *n* = 0 and *c* ∈ [10^−4^, 10^−2^]M for *n* = 4) the ion channel spends at least 10% of its time in the remaining states which results in fundamentally different dynamics than what is predicted by the Hill response and (2) even in the limits such as *c* = 0 and *c* → ∞ where the *C*_0_ and *O*_4_ states dominate the system, the Hill limit ignores the leakiness and dynamic range of the ion channel (requiring them to exactly equal 0 and 1, respectively), thereby glossing over these important properties of the system.

**Figure 7:**
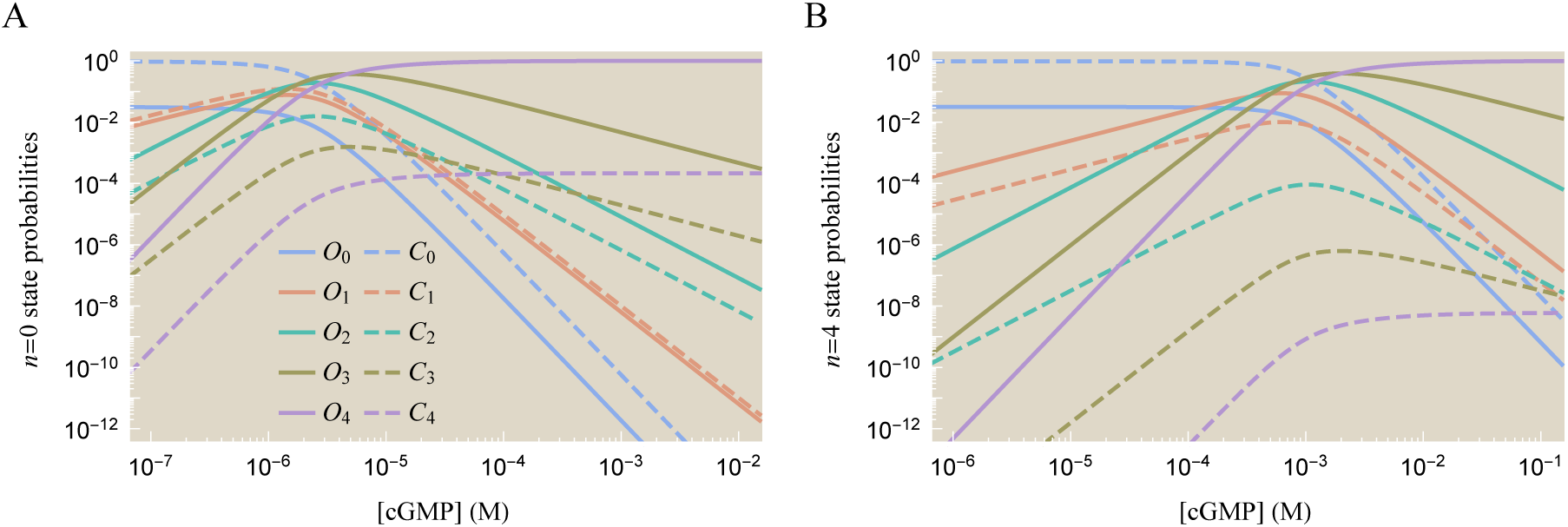
Individual state probabilities for the wild type and mutant CNGA2 ion channels. (A) The state probabilities for the wild type (*n* = 0) ion channel. The subscripts of the open (*O_j_*) and closed (*C_j_*) states represent the number of ligands bound to the channel. States with partial occupancy, 1 ≤ *j* ≤ 3, are most likely to occur in a narrow range of ligand concentrations [cGMP] ∈ [10^−7^, 10^−5^] M, outside of which either the completely empty *C*_0_ or fully occupied *O*_4_ states dominate the system. (B) The state probabilities for the *n* = 4 channel. Because the mutant subunits have a weaker affinity to ligand (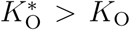 and 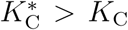), the state probabilities are all shifted to the right.

### Characterizing CNGA2 Mutants based on Subunit Composition

We now turn to the leakiness, dynamic range, [*EC*_50_], and effective Hill coefficient *h* of a CNGA2 ion channel with *n* mutated and *m* − *n* wild type subunits. Detailed derivations for the following results are provided in Supporting Information section B:

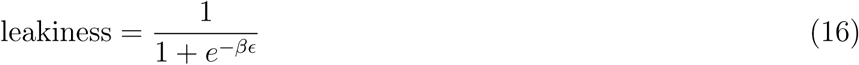

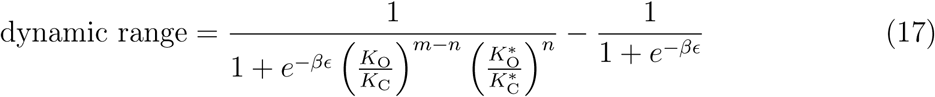

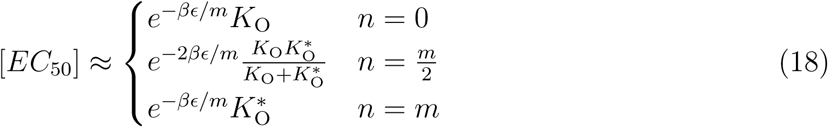

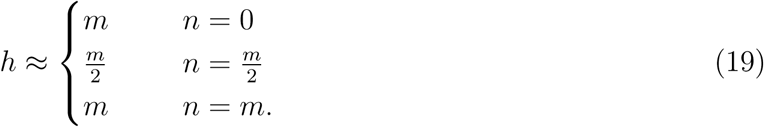

Note that we recover the original MWC model results Eqs (5)-(12) for the *n* = 0 wild type ion channel. Similarly, the homooligomeric *n* = *m* channel is also governed by the MWC model with 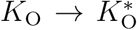 and 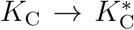. We also show the [*EC*_50_] and *h* formulas for the 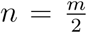 case to demonstrate the fundamentally different scaling behavior that this heterooligomeric channel exhibits with the MWC parameters.

Fig 8A shows that all of the CNGA2 mutants have small leakiness, which can be understood from their small *ϵ* value and Eq (16). In addition, the first term in the dynamic range Eq (17) is approximately 1 because the open state affinities are always smaller than the closed state affinities by at least a factor of ten, which is then raised to the fourth power. Thus, all of the mutants have a large dynamic range as shown in Fig 8B. Experimentally, single channel measurements confirmed that the wild type *n* = 0 channel is nearly always closed in the absence of ligand; in the opposite limit of saturating cGMP, it was found that 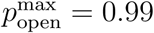 for both the *n* = 0 and *n* = *m* ion channels (see Supporting Information section C.2).^13^

**Figure 8:**
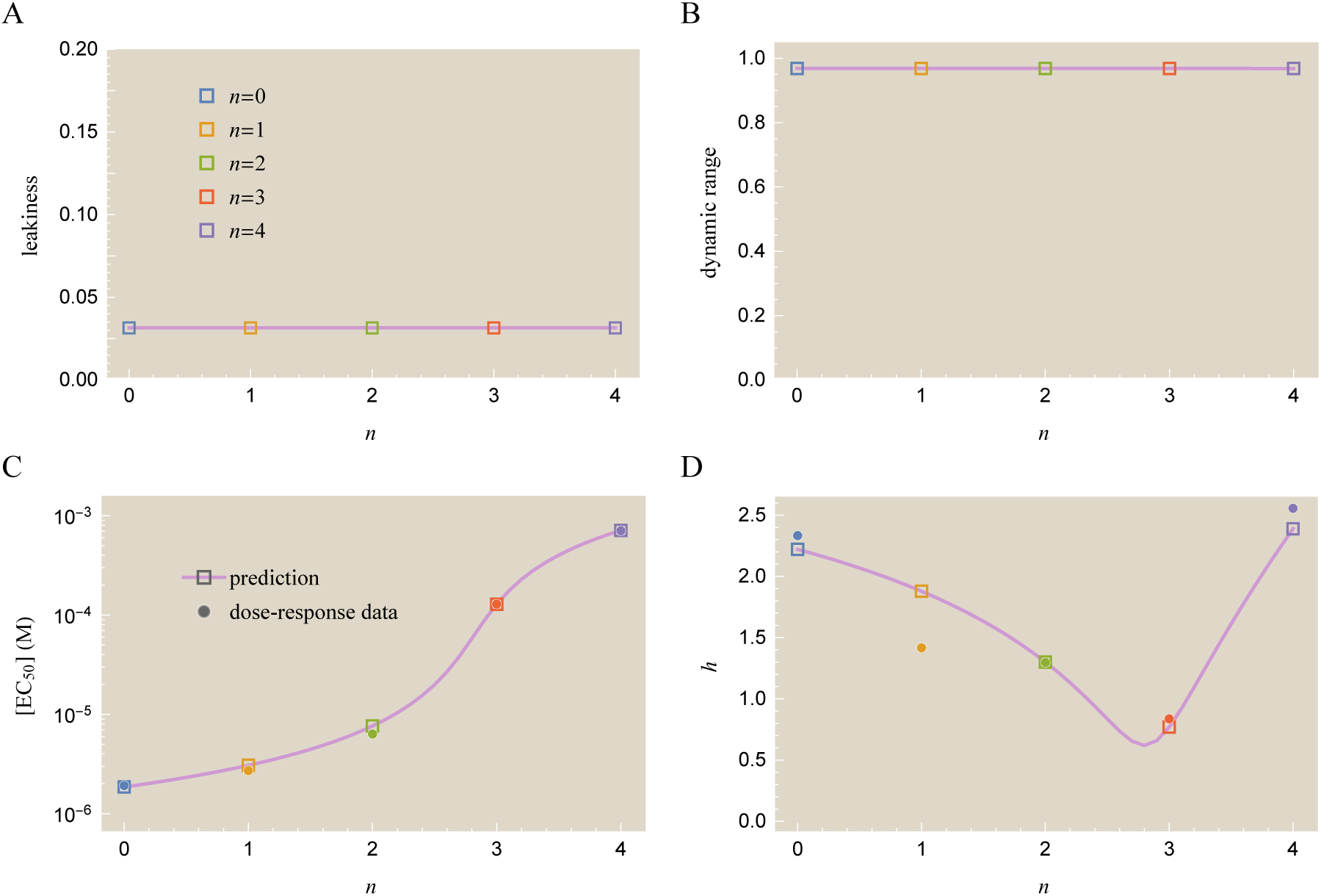
Theoretical prediction and experimental measurements for mutant CNGA2 ion channel characteristics. The open squares represent the five mutant ion channels in Fig 6 with *n* mutated subunits. (A) All ion channels have small leakiness. (B) The dynamic range of all channels is near the maximum possible value of unity, indicating that they rarely open in the absence of ligand and are always open in the presence of saturating ligand concentrations. (C) The [*EC*_50_] increases non-uniformly with the number of mutant subunits. Also shown are the measured values (solid circles) interpolated from the data. (D) The effective Hill coefficient has a valley due to the competing influences of the wild type subunits (which respond at *μ*M ligand concentrations) and the mutant subunits (which respond at mM concentrations). Although the homotetrameric channels (*n* = 0 and *n* = 4) both have sharp responses, the combined effect of having both types of subunits (*n* = 1, 2, and 3) leads to a flatter response.

The [*EC*_50_] and effective Hill coefficient *h* are shown in Fig 8C and D. In contrast to the nAChR case, where each additional mutation decreased [*EC*_50_], each CNGA2 mutation tends to increase [*EC*_50_], although not by a uniform amount. The effective Hill coefficient has a particularly complex behavior, first decreasing with each of the first three subunit mutations and then finally increasing back to the wild type level for the fully mutated ion channel. To explain this decrease in *h* for the heterooligomeric channels, we first note that the wild type *n* = 0 channel has a sharp response about its [*EC*_50_] ≈ *e*^*−βϵ*/*m*^*K*_O_ while the fully mutated *n* = *m* channel has a sharp response about 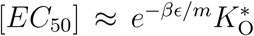. Roughly speaking, the response of the heterooligomeric channels with 1 ≤ *n* ≤ 3 will occur throughout the full range between *e*^*−βϵ*/*m*^*K*_O_ and 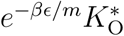, which causes the dose-response curves to flatten out and leads to the smaller effective Hill coefficient. Such behavior could influence, for example, the response of the heterooligomeric nAChR ion channel if the two acetylcholine binding pockets diverged to have different ligand affinities.

Although we have focused on the particular mutants created by Wongsamitkul *et al.*, it is straightforward to apply this framework to other types of mutations. For example, in Supporting Information section B.2 we consider the effect of modifying the *K*_O_ and *K*_C_ parameters of all four CNGA2 channels simultaneously. This question is relevant for physiological CNGA2 channels where a mutation in the gene would impact all of the subunits in the homooligomer, in contrast to the Wongsamitkul constructs where the four subunits were stitched together within a single gene. We find that when *K*_O_ and *K*_C_ vary for all subunits, the leakiness, dynamic range, and effective Hill coefficient remain nearly fixed for all parameter values, and that only the [*EC*_50_] scales linearly with *K*_O_ as per Eq (11). In order to affect the other properties, either the gating energy *βϵ* or the number of subunits *m* would need to be changed.

### Extrapolating the Behavior of a Class of Mutants

In this section, we explore how constant trends in both the nAChR and CNGA2 data presented above provide an opportunity to characterize the full class of mutants based on the dose-response curves from only a few of its members. Such trends may well be applicable to other ion channel systems, enabling us to theoretically probe a larger space of mutants than what is available from current data.

First, we note that because the [*EC*_50_] of the five nAChR mutants fell on a line in Fig 4C, we can predict the response of the entire class of mutants by only considering the dose-response curves of two of its members and extrapolating the behavior of the remaining mutants using linear regression. Experimentally, such a characterization arises because the L251S mutation acts nearly identically and independently across subunits to change the gating free energy of nAChR.^12,17,23^ This implies that mutating *n* subunits would yield an ion channel with gating energy

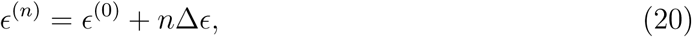

where *ϵ*^(0)^ is the wild type gating energy and Δ*ϵ* is the change in free energy per mutation. This functional form is identical to the mismatch model for transcription factor binding, where each additional mutation-induced mismatch adds a constant energy cost to binding.^27^ Fig 9A demonstrates how fitting only the *n* = 0 and *n* = 4 dose-response curves (solid lines) accurately predicts the behavior of the *n* = 1, 2, and 3 mutants (dashed lines). In Supporting Information section D, we carry out such predictions using all possible input pairs.

**Figure 9:**
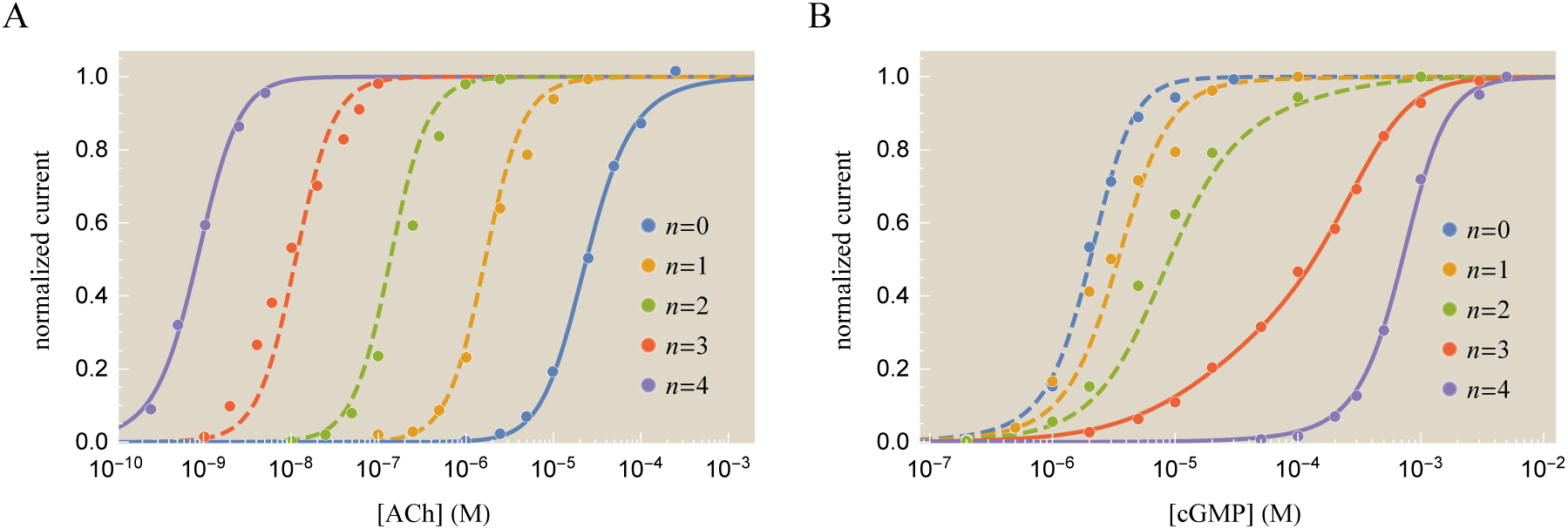
Predicting the dose-response of a class of mutants using a subset of its members. (A) The MWC parameters of the nAChR mutants can be fixed using only two data sets (solid lines), which together with Eq (20) predict the dose-response curves of the remaining mutants (dashed lines). (B) For the CNGA2 channel, the properties of both the wild type and mutant subunits can also be fit using two data sets, accurately predicting the responses of the remaining three mutants. Supporting Information section D demonstrates the results of using alternative pairs of mutants to fix the thermodynamic parameters in both systems.

We now turn to the CNGA2 ion channel where, once the *K*_O_, *K*_C_, 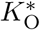, 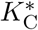, and *ϵ* parameters are known, the dose-response curve of any mutant can be predicted by varying *n* in Eq (15). Fig 9B demonstrates that the wild type ion channel (*n* = 0) and the ion channel with only one mutated subunit (*n* = 1) can accurately predict the dose-response curves of the remaining mutants. Supporting Information section D explores the resulting predictions using all possible input pairs.

### MWC Model allows for Degenerate Parameter Sets

One critical aspect of extracting model parameters from data is that degenerate sets of parameters may yield identical outputs, which suggests that there are fundamental limits to how well a single data set can fix parameter values.^25,28^ This phenomenon, sometimes dubbed “sloppiness,” may even be present in models with very few parameters such as the MWC model considered here. Fig 10 demonstrates the relationship between the best-fit parameters within the nAChR and CNGA2 systems. For concreteness, we focus solely on the nAChR system.

**Figure 10:**
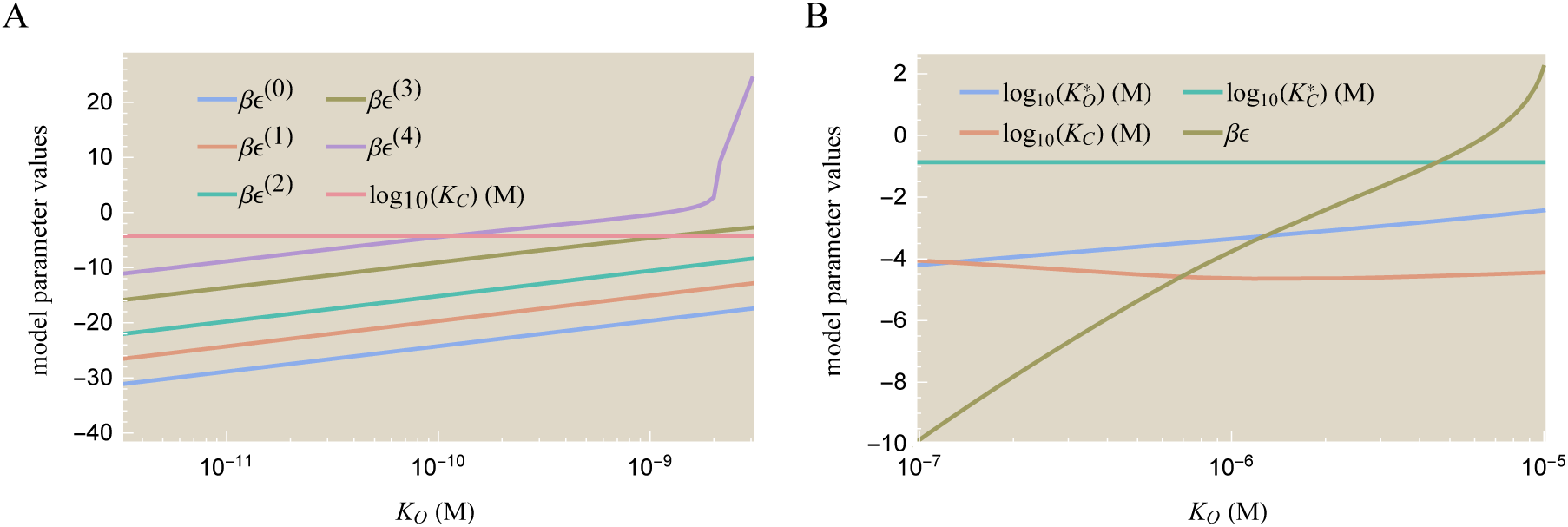
Degenerate parameter sets for nAChR and CNGA2 model fitting. Different sets of biophysical parameters can yield the same system response. (A) Data for the nAChR system in Fig 3 is fit by constraining *K*_O_ to the value shown on the x-axis. The remaining parameters can compensate for this wide range of *K*_O_ values. (B) The CNGA2 system in Fig 6 can similarly be fit by constraining the *K*_O_ value, although fit quality decreases markedly outside the narrow range shown. Any set of parameters shown for either system leads to responses with *R*^2^ > 0.96.

After fixing the value of *K*_O_ (to that shown on the x-axis of Fig 10A), the remaining parameters are allowed to freely vary in order to best fit the nAChR data. Although every value of *K*_O_ ϵ [10^−11^ − 10^−9^]M yields a nearly identical response curve in excellent agreement with the data (with a coefficient of determination *R*^2^ > 0.96), we stress that dissociation constants are rarely found in the range *K*_O_ ≪ 10^−10^ M. In addition, a dissociation constant above the nM range, *K*_O_ ≫ 10^−9^ M, cannot fit the data well and is therefore invalidated by the model. Thus, we may suspect the true parameter values will fall around the interval *K*_O_ ϵ [10^−10^ − 10^−9^]M for the nAChR system. *K*_O_ could ultimately be fixed by measuring the leakiness Eq (5) (and thereby fixing *βϵ*) for any of the ion channel mutants.

Two clear patterns emerge from Fig 10A: (1) The value of *K*_C_ is approximately constant for all values of *K*_O_ and (2) the five free energies all vary as *βϵ*^(n)^ = 2 log (*K*_O_)+*n* (constant). This suggests that *K*_C_ and *e*^*−βϵ*/2^*K*_O_ are the fundamental parameters combinations of the system. In Supporting Information section C.4, we discuss how these parameter combinations arise from our model.

We end by noting that the notion of sloppiness, while detrimental to fixing the physical parameter values of a system, nevertheless suggests that multiple evolutionary paths may lead to optimal ion channel phenotypes, providing another mechanism by which allostery promotes a protein’s capacity to adapt.^29^

## Discussion

There is a deep tension between the great diversity of biological systems and the search for unifying perspectives to help integrate the vast data that has built up around this diversity. Several years ago at the Institut Pasteur, a meeting was convened to celebrate the 50^th^ anniversary of the allostery concept, which was pioneered in a number of wonderful papers in the 1960s and which since then has been applied to numerous biological settings.^30–34^ Indeed, that meeting brought together researchers working in physiology, neuroscience, gene regulation, cell motility, and beyond, making it very clear that allostery has great reach as a conceptual framework for thinking about many of the key macromolecules that drive diverse biological processes.

In this paper, we have built upon this significant previous work and explored how the Monod-Wyman-Changeux model can serve as a unifying biophysical framework for whole suites of ion channel mutants (see Figs 3 and 6). Specifically, we used two well-studied ligand-gated ion channels to explore the connection between mutations, the MWC parameters, and the full spectrum of dose-response curves which may be induced by those mutations. In addition, we have shown how earlier insights into the nature of “data collapse” in the context of bacterial chemotaxis and quorum sensing^19,20^ can be translated into the realm of ion channels. By introducing the Bohr parameter, we are able to capture the non-linear combination of thermodynamic parameters which governs the system’s response.

For both the nAChR and CNGA2 ion channels, we showed that precise predictions of dose-response curves can be made for an entire class of mutants by only using data from two members of this class (Fig 9). In other words, the information contained in a single dose-response curve goes beyond merely providing data for that specific ion channel. Ultimately, because the total space of all possible mutants is too enormous for any significant fraction to be explored experimentally, we hope that a coupling of theory with experiment will provide a step towards mapping the relation between channel function (phenotype) and the vast space of protein mutations.

Moreover, we used the MWC model to determine analytic formulas for key properties such as the leakiness, dynamic range, [*EC*_50_], and the effective Hill coefficient, which together encapsulate much of the information in dose-response curves. These relationships tie into the extensive knowledge about phenotype-genotype maps,^27,29,35^ enabling us to quantify the trade-offs inherent in an ion channel’s response. For example, when modifying the ion channel gating free energy, the changes in the leakiness and [*EC*_50_] are always negatively correlated (Fig 4), whereas modifying the ligand binding domain will not affect the leakiness but may change the [*EC*_50_] (Fig 8 and Supporting Information section B.2). The ability to navigate between the genotype and phenotype of proteins is crucial in many bioengineering settings, where site-directed mutagenesis is routinely employed to find mutant proteins with specific characteristics (e.g. a low leakiness and large dynamic range).^36–38^

While general formulas for these phenotypic properties were elegantly derived in earlier work,10 we have shown that such relations can be significantly simplified in the context of ion channels where 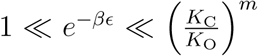 (see Eqs (9)-(12)). This approximation is applicable for the range of parameters spanned by both the nAChR and CNGA2 systems, and we suspect it may hold for many other ion channels. These formulas provide a simple, intuitive framework to understand the effects of mutations. For example, they suggest: (1) Channel pore mutations that increase *ϵ* will exponentially increase the leakiness of the channel, although the constraint 1 ≪ *e*^−*βϵ*^ ensures that this leakiness will still be small. Ligand domain mutations are not expected to affect leakiness at all. (2) Channel pore mutations will exponentially decrease the [*EC*_50_] with increasing *ϵ*, although this effect is diminished for ion channels with multiple subunits. For mutations in the ligand binding domain, the [*EC*_50_] will increase linearly with the dissociation constant *K*_O_ between the ligand and the open ion channel (see Supporting Information section B.2). (3) Neither the dynamic range nor the effective Hill coefficient will be significantly perturbed by either type of mutation. (4) Transforming a homooligomeric ion channel into a heterooligomer can generate a significantly flatter response. For example, even though the CNGA2 channel comprised of either all wild type or all mutant subunits had a very sharp response, a channel comprised of both subunits had a smaller effective Hill coefficient (see Fig 8D).

The framework presented here could be expanded in several exciting ways. First, it remains to be seen whether channel pore mutations and ligand binding domain mutations are completely independent, or whether there is some cross-talk between them. This question could be probed by creating a channel pore mutant (whose dose-response curves would fix its new 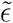 values), a ligand domain mutant (whose new 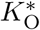 and 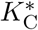 values would be characterized from its response curve), and then creating the ion channel with both mutations. If these two mutations are independent, the response of the double mutant can be predicted *a priori* using 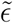, 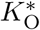, and 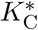.

We also note that the MWC model discussed here does not consider several important aspects relating to the dynamics of ion channel responses. Of particular importance is the phenomenon of desensitization which significantly modifies an ion channel’s response in physiological settings.^39,40^ In addition, some ion channels have multiple open and closed conformations^41–43^ while other channels exhibit slow switching between the channel states.^44^ Exploring these additional complexities within generalizations of the MWC model would be of great interest.

Finally, we believe that the time is ripe to construct an explicit biophysical model of fitness to calculate the relative importance of mutation, selection, and drift in dictating the diversity of allosteric proteins such as the ion channels considered here. Such a model would follow in the conceptual footsteps laid in the context of fitness of transcription factors binding,^27,35,45^ protein folding stability,^46–48^ and influenza evolution.^49^ This framework would enable us to make precise, quantitative statements about intriguing trends; for example, nearly all nAChR pore mutations appear to increase a channel’s leakiness, suggesting that minimizing leakiness may increase fitness.12 One could imagine that computing derivatives such as 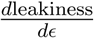, a quantity analogous to the magnetic susceptibility in physics, would be correlated with how likely an ϵ mutation is to be fixed. The goal of such fitness functions is to map the complexity of the full evolutionary space (i.e. changes to a protein amino acid sequence) onto the MWC parameters, and then the fitness function determines how these parameters evolve in time. In this way, the complexity of sequence and structure would fall onto the very low dimensional space governed by *ϵ*, *K*_O_, and *K*_C_.

## Acknowledgement

It is with sadness that we dedicate this paper to the memory of Klaus Schulten with whom one of us (RP) wrote his first paper in biophysics. Klaus was an extremely open and kind man, a broad and deep thinker who will be deeply missed. We thank Stephanie Barnes, Nathan Belliveau, Chico Camargo, Griffin Chure, Lea Goentoro, Michael Manhart, Chris Miller, Muir Morrison, Manuel Razo-Mejia, Noah Olsman, Allyson Sgro, and Jorge Zañudo for their sharp insights and valuable feedback on this work. We are also grateful to Henry Lester, Klaus Benndorf, and Vasilica Nache for helpful discussions as well as sharing their ion channel data. All plots were made in *Mathematica* using the CustomTicks package.^50^ This work was supported by La Fondation Pierre-Gilles de Gennes, the Rosen Center at Caltech, the National Science Foundation under NSF PHY11-25915 at the Kavli Center for Theoretical Physics, and the National Institutes of Health through DP1 OD000217 (Director’s Pioneer Award), 5R01 GM084211C, R01 GM085286, and 1R35 GM118043-01 (MIRA).

### Supporting Information Available

The following Supporting Information is available:

- Details on aforementioned derivations and calculations (PDF)
- All of the plots in the main text can be reproduced with the supplementary *Mathematica* notebook. The dose-response data for both the nAChR and CNGA2 systems may be found in this notebook (ZIP)

This material is available free of charge via the Internet at http://pubs.acs.org/.

